# Sequencing and variant detection of eight abundant plant-infecting tobamoviruses across Southern California wastewater

**DOI:** 10.1101/2022.08.03.502731

**Authors:** Jason A. Rothman, Katrine L. Whiteson

**Author notes:** Corresponding author: Jason A. Rothman, University of California, Irvine, Irvine, CA, 92697.

## Abstract

Tobamoviruses are agriculturally-relevant viruses that cause crop losses and have infected plants in many regions of the world. These viruses are frequently found in municipal wastewater - likely coming from human diet and industrial waste across wastewater catchment areas. As part of a large wastewater-based epidemiology study across Southern California, we analyzed RNA sequence data from 275 influent wastewater samples obtained from eight wastewater treatment plants with a catchment area of approximately 16 million people from July 2020 – August 2021. We assembled 1,083 high-quality genomes, enumerated viral sequencing reads, and detected thousands of single nucleotide variants from eight common tobamoviruses: Bell Pepper Mottle Virus, Cucumber Green Mottle Mosaic Virus, Pepper Mild Mottle Virus, Tobacco Mild Green Mosaic Virus, Tomato Brown Rugose Fruit Virus, Tomato Mosaic Virus, Tomato Mottle Mosaic Virus, and Tropical Soda Apple Mosaic Virus. We show that single nucleotide variants had amino acid-altering consequences along with synonymous mutations, which represents potential evolution with functional consequences in genomes of these viruses. Our study shows the importance of wastewater sequencing to monitor the genomic diversity of these plant-infecting viruses, and we suggest that our data could be used to continue tracking the genomic variability of such pathogens.

**Importance:** Diseases caused by viruses in the genus *Tobamovirus* cause crop losses around the world. As with other viruses, mutation occurring in the virus’ genomes can have functional consequences and may alter viral infectivity. Many of these plant-infecting viruses have been found in wastewater, likely coming from human consumption of infected plants and produce. By sequencing RNA extracted from influent wastewater obtained from eight wastewater treatment plants in Southern California, we assembled high-quality viral genomes and detected thousands of single nucleotide variants from eight tobamoviruses. Our study shows that *Tobamovirus* genomes vary at many positions, which may have important consequences to viral host switching and the detection of these viruses by agricultural or environmental scientists.

## Introduction

Wastewater represents a matrix of microorganisms, human waste, and water inflow across a sewage catchment area (1). As part of the microorganismal fraction of wastewater, there are often high abundances of plant-infecting +ssRNA viruses of the genus *Tobamovirus* - which represents important plant pathogens causing substantial crop losses to the agricultural industry (2–6). Tobamoviruses are widespread and may be deposited into wastewater through agricultural runoff and human diet, where they can resist degradation even through wastewater and drinking water treatment (2). They are often the most abundant RNA viruses in human feces and wastewater samples, even going back to the first human fecal RNA virome sequenced (7). As part of ongoing efforts and advances in wastewater-based epidemiology (WBE), it is critical to monitor wastewater for the presence of tobamoviruses and their potential to infect new hosts or evade plant immunity (8). Also, as many tobamoviruses may serve as water quality indicators and are impactful diseases to agriculture, studies should be conducted to understand the genomics of these viruses (2, 9).

As part of a large WBE effort across Southern California, we used metatranscriptomic sequencing to investigate the genomics and single nucleotide variants (SNVs) of eight tobamoviruses sourced from 275 samples across eight wastewater treatment plants from July 2020 – August 2021 (3, 10). These viruses were Bell Pepper Mottle Virus (BPeMV), Cucumber Green Mottle Mosaic Virus (CGMMV), Pepper Mild Mottle Virus (PMMoV) Tobacco Mild Green Mosaic Virus (TMGMV), Tomato Brown Rugose Fruit Virus (ToBRFV), Tomato Mosaic Virus (ToMV), Tomato Mottle Mosaic Virus (ToMMV), and Tropical Soda Apple Mosaic Virus (TSAMV). Through our study, we investigated several lines of inquiry: Can we assemble high-quality *Tobamovirus* genomes from wastewater samples? Do we obtain acceptable sequencing coverage across viral genomes derived from wastewater? Can we identify SNVs across tobamoviruses in Southern California’s wastewater?

## Materials and Methods

We obtained raw sequencing data as FASTQ files from the NCBI Sequence Read Archive under BioProject PRJNA729801, and we refer to Rothman, et al 2021 (3) and Rothman et al 2022 (10) for all sampling, RNA extraction, and sequencing methods. We used BBTools (11) “bbduk” to remove sequencing adapters, primers, and low-quality bases from the reads, BBTools “dedupe” to remove optical duplicates, and removed human genome reads (hg38) with Bowtie2 (12). We then used Bowtie2 to align the reads to the reference strains (downloaded from NCBI) for each *Tobamovirus*: BPeMV (NC_009642.1), CGMMV (NC_001801.1), PMMoV (NC_003630.1), TMGMV (NC_001556.1), ToBRFV (NC_028478.1), ToMV (NC_002692.1), ToMMV (NC_022230.1), and TSAMV (NC_030229.1).

We used samtools (13) to assess sequencing depth and breadth of genomic coverage on the BAM files. We then used iVar (14) to identify single nucleotide variants (SNVs) for each virus in each sample separately and plotted the SNVs and genome depth/coverage in R (15) using “ggplot2” (16) and “patchwork” (17). We assembled contigs within each sample with MEGAHIT (18) and assessed contig assembly quality with checkV (19), using a cutoff of > 90% completeness and 0% contamination to characterize them as “high-quality genomes.” We used DIAMOND (20) to classify the “high-quality genomes” and VIRIDIC (21) to calculate the intergenomic similarity between viral genomes, and plotted summary statistics about each sample and virus with “ggplot2.”

Data used in this study are available on the NCBI Sequence Read Archive under accession # PRJNA729801 and on the Dryad Digital Repository (doi.org/10.7280/D1S69X) (22).

## Results

We aligned 156,825,269 quality-filtered, deduplicated, matching paired-end reads (313,650,538 individual reads) across 275 samples from eight water treatment plants (average = 570,274 paired-end reads, range = 44 – 8,933,433). Of the paired-end reads that mapped to the eight tobamoviruses, 0.34% were Bell Pepper Mottle Virus (BPeMV), 12.90% were Cucumber Green Mottle Mosaic Virus (CGMMV), 11.90% were Pepper Mild Mottle Virus (PMMoV), 1.31% were Tobacco Mild Green Mosaic Virus (TMGMV), 64.06% were Tomato Brown Rugose Fruit Virus (ToBRFV), 5.6% were Tomato Mosaic Virus (ToMV), 1.87% were Tomato Mottle Mosaic Virus (ToMMV), and 2.02% were Tropical Soda Apple Mosaic Virus (TSAMV) (Fig. 1, Fig. S1).

**Figure 1:**
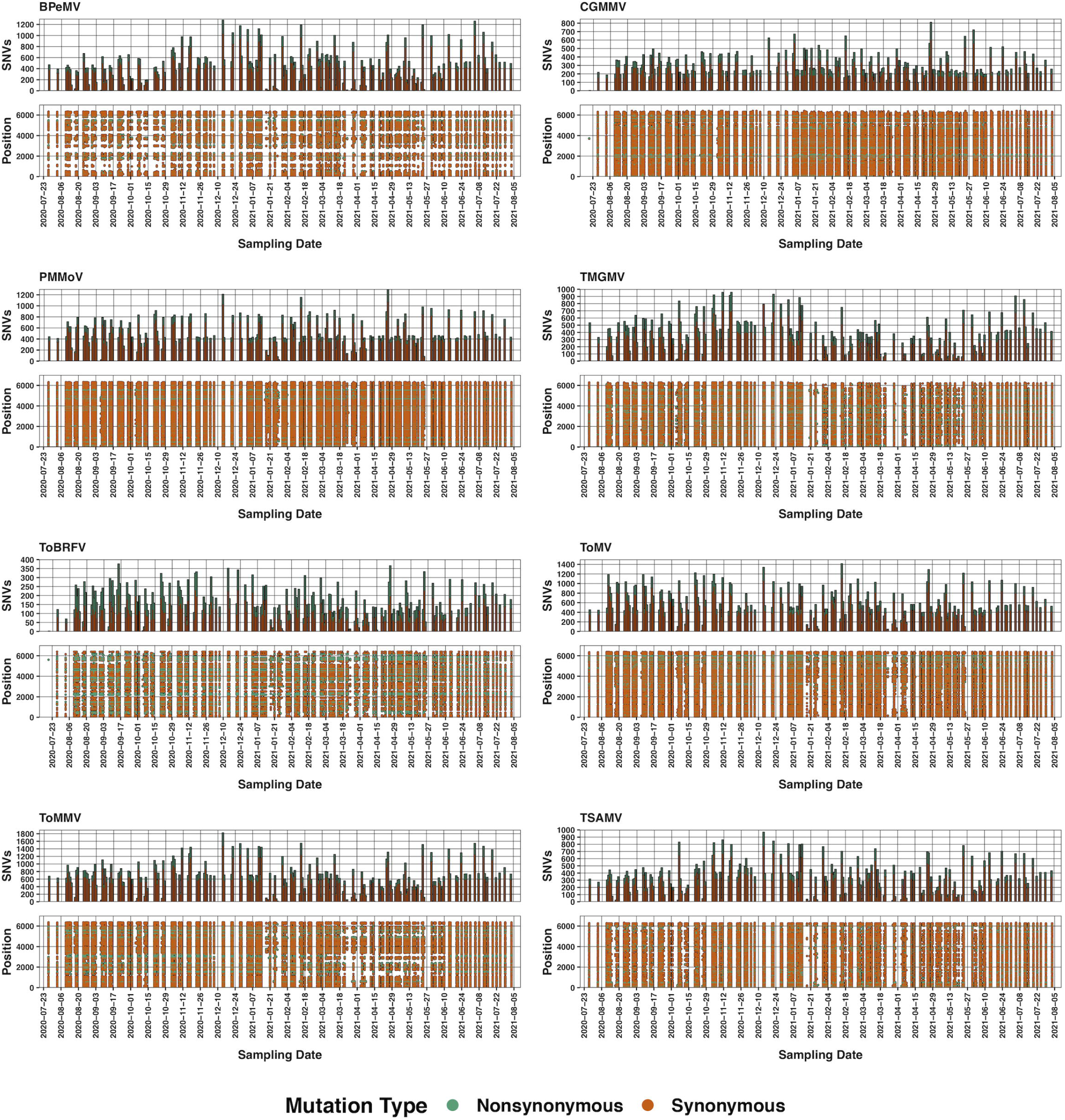
Boxplots of the average relative abundances of mapped reads (within this study only) of each *Tobamovirus* across all samples. Lines within each box represents the median relative abundance, whiskers are 1.5× IQR, and dots are values >1.5 IQR.

For each virus, we report the total number, average, and range of mapped paired-end reads, the average sequencing depth and overall genomic coverage, number of high-quality assembled genomes, average genomic similarity between assembled genomes as determined by VIRIDIC, the minimum DIAMOND alignment percentage, and the number of single nucleotide variants (SNVs) along with the SNVs’ mutational consequence (synonymous or nonsynonymous) in Table 1. We also plotted the average read depth per base pair (Fig. S2), and the genomic position and date of each SNV detected along with its mutational consequence for each virus (Fig. 2). Lastly, we provide the relevant iVAR output for each sample and SNV along with the sequences of all high-quality viral genomes on Dryad (doi.org/10.7280/D1S69X)(22).

**Table 1:** Sequencing results for each virus across all viruses.

| Virus | Mapped Paired-End Reads | Average/Range Mapped Reads Per Sample | Average Sequencing Depth Per Base and Breadth | Number of High-Quality Genomes | Average Genomic Similarity Between Genomes | Alignment Percentage of Genomes to Reference Strain | SNVs |
| --- | --- | --- | --- | --- | --- | --- | --- |
| BPeMV | 526,713 | 1,915<br>(0 - 26,075) | 84x<br>99.5% | 0 | N/A | N/A | 963:<br>dN = 241, dS = 722 |
| CGMMV | 20,222,039 | 73,535<br>(6 - 1,248,760) | 2,347x<br>100% | 144 | 99.6% | >99.5% | 1,384:<br>dN = 397, dS = 987 |
| PMMoV | 18,663,282 | 67,866<br>(0 - 1,028,539) | 2,130x<br>100% | 90 | 98.4% | >99.3% | 1,306:<br>dN = 381, dS = 925 |
| TMGMV | 2,057,301 | 7,481<br>(0 - 116,449) | 240x<br>100% | 141 | 99.2% | >97.6% | 1,557:<br>dN = 509, dS = 1,048 |
| ToBRFV | 100,455,804 | 365,294<br>(1 - 6,727,467) | 11,854x<br>100% | 250 | 99.7% | >99.9% | 1,075:<br>dN = 452, dS = 623 |
| ToMV | 8,788,061 | 31,957<br>(0 - 513,129) | 1,162x<br>99.9% | 183 | 98.8% | >99.6% | 1,540:<br>dN = 429, dS = 1,111 |
| ToMMV | 2,939,466 | 10,689 (0 - 120,789) | 487x<br>99.9% | 79 | 98.5% | >98.6% | 1,309:<br>dN = 372, dS = 937 |
| TSAMV | 3,172,603 | 11537 (0 - 200383) | 360x<br>100% | 196 | 99.3% | >99.4% | 1,531:<br>dN = 486, dS = 1045 |

**Figure 2:**
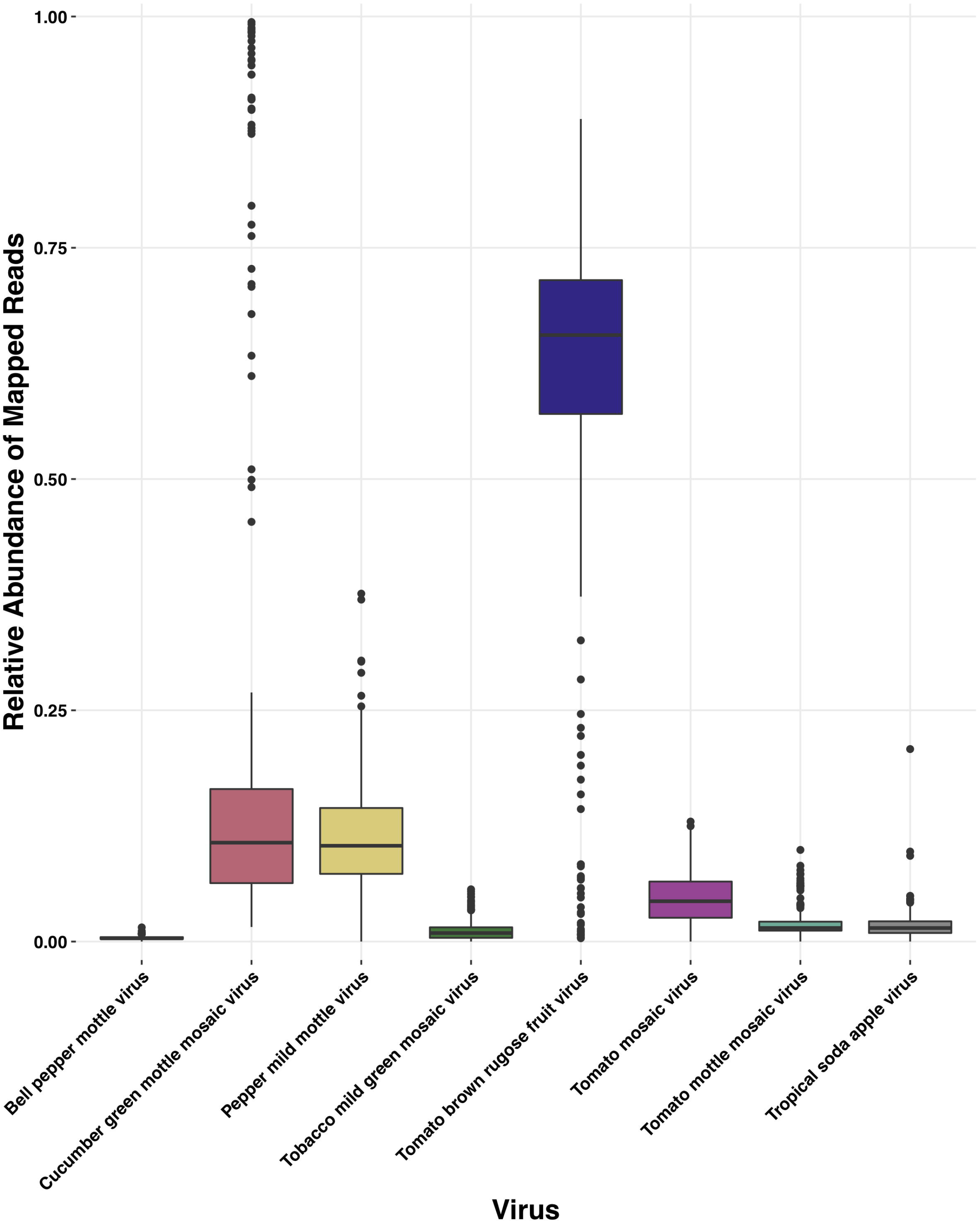
Genomic position and number of single nucleotide variants (SNVs) for BPeMV, CGMMV, PMMoV, TMGMV, ToBRFV, ToMV, ToMMV, and TSAMV at each sampling date. Plot coloration denotes mutational consequence of the SNV.

## Discussion

Wastewater-based epidemiology (WBE) has been used to characterize pathogen abundances and genomics for a variety of diseases and is often employed to detect antibiotic resistance or diseases relevant to public health (5). We applied similar molecular and bioinformatic methods to eight agriculturally-relevant tobamoviruses sequenced from influent wastewater, representing a sewer shed of approximately 16 million Southern Californians across eight wastewater treatment plants (3, 10). These tobamoviruses were abundant and widespread throughout our wastewater samples, comprising eight of the top 10 viruses in our dataset (10). PMMoV may be the best known and is often regarded as the most abundant virus in fecal and wastewater samples (7, 9), however, we were surprised to find that ToBRFV was much more abundant, mirroring the results of a recent study from Maryland (5). Likely due to their near-ubiquity, we obtained very deep and broad sequencing coverage across their genomes. Our samples yielded thousands of SNVs per virus, and we assembled over 70 individual high-quality genomes for each viral species except BePMV, supporting studies that suggest WBE is useful to characterizing the genomic landscape of pathogens (3, 5, 6, 23). Interestingly, most of the SNVs identified were synonymous mutations, although there were thousands of putative nonsynonymous mutations that may have consequences in host infectivity or immune escape.

As tobamoviruses are being developed for use as water quality indicators, it is important to have a broad pool of wastewater-sourced genomes, sequences, and SNVs so that proper tests can be developed that reflect the diversity of each virus (2, 9). For example, *Tobamovirus* testing involves careful selection of specific, validated RT-qPCR primers, which may lose specificity as viral mutations arise, making pathogen detection unreliable without adjusting for new variants (24). Likewise, to combat outbreaks of tobamoviruses, host-switching, or the evolution of novel viruses, deep sequence resources should be provided to the scientific and agricultural communities (8). For ToBRFV - the most abundant *Tobamovirus* in our dataset - intergenomic similarity was >97% across all genomes, making ToBRFV a good target for RT-qPCR assays, and an important water quality indicator in an era when water reuse is becoming more critical (9). To the best of our knowledge, our study is the first to sequence such a wide diversity of *Tobamovirus* genomes and SNVs from wastewater, and we suggest that future research be conducted using WBE for other agriculturally-relevant diseases. Furthermore, as water reuse is becoming widespread, studies should investigate the ability of wastewater treatment plants to inactivate tobamoviruses to prevent accidental infection through irrigation, and to indicate expected decreases in viral load for public health (9).

## Acknowledgments

This research was supported by the University of California Office of the President Research Grants Program Office (award numbers R01RG3732 and R00RG2814) awarded to JAR and KLW, and a Hewitt Foundation for Biomedical Research postdoctoral fellowship to JAR. This work was made possible, in part, through access to computing resources from the UCI High Performance Community Computing Cluster and sequencing assistance of the UCI Genomics High-Throughput Facility. We thank Susan Hiestand and Eric Martens for the interesting conversation about the observation of uncultivated tomatoes growing along wastewater streams and we thank the Southern California Coastal Water Research Project for fruitful collaborations and thoughtful conversations.

