## Supplemental information for "Sequencing and variant detection of eight abundant plant-infecting tobamoviruses across Southern California wastewater"

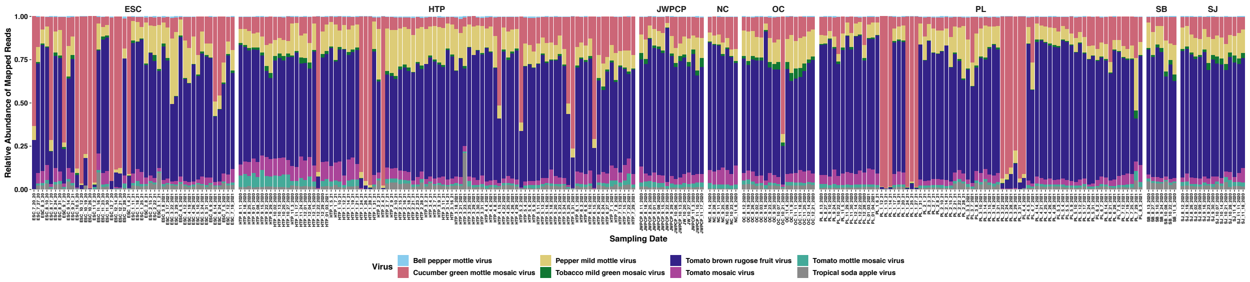


Figure S1: Stacked bar plot of the relative abundances of mapped paired-end reads for each *Tobamovirus* in individual samples within this study, faceted by water treatment plant. Colors denote virus identity.

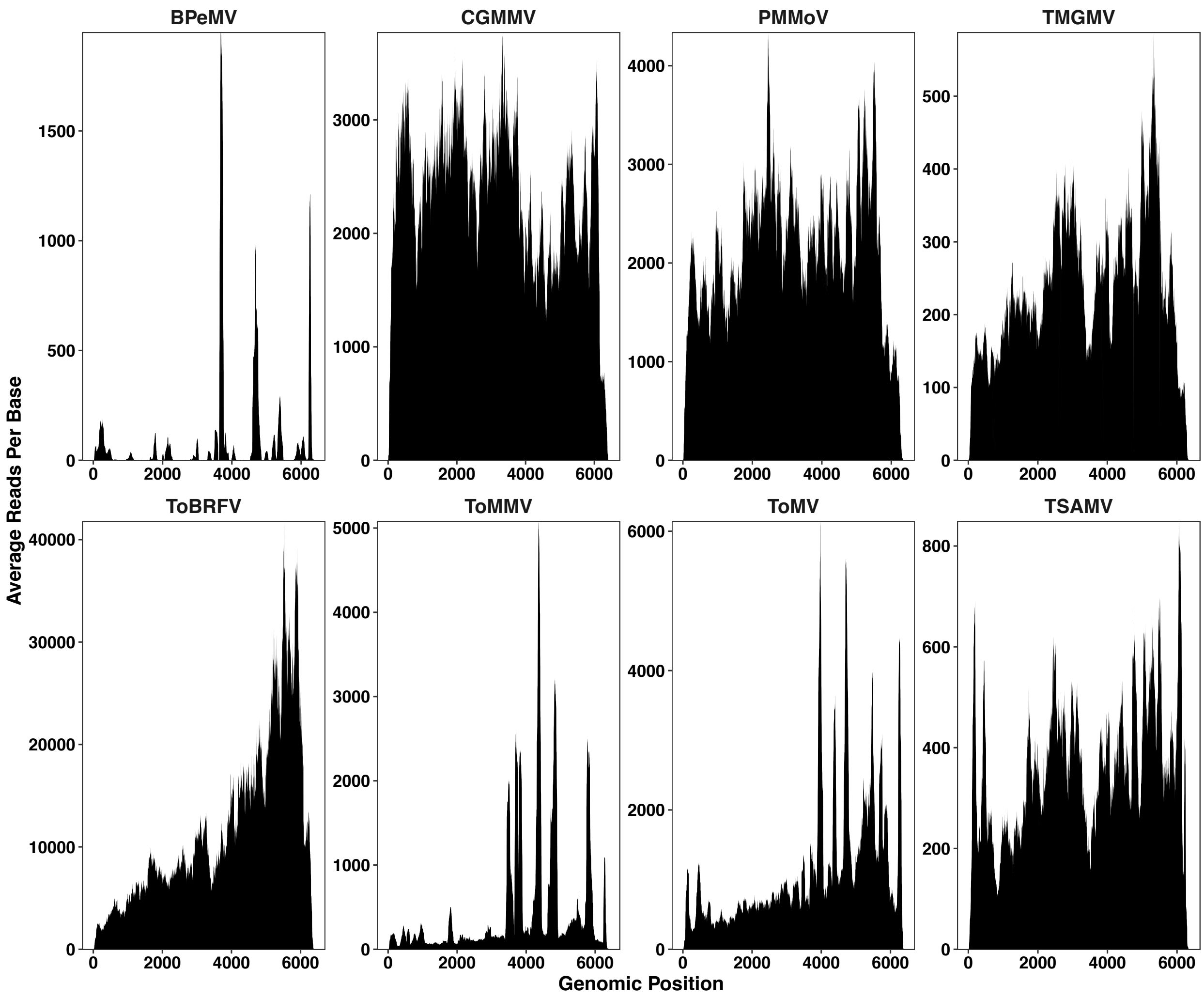


Figure S2: Area plot of the average paired-end mapped reads per base across all samples for each *Tobamovirus* faceted by virus. X-axes represent the genomic nucleotide position within each virus.
